# A pairwise maximum entropy model uncovers the white matter scaffold underlying emergent dynamics in intracranial EEG

**DOI:** 10.1101/507962

**Authors:** Arian Ashourvan, Preya Shah, Adam Pines, Shi Gu, Christopher W. Lynn, Danielle S. Bassett, Kathryn A. Davis, Brian Litt

## Abstract

A major challenge in systems neuroscience is to understand how the brain’s structural architecture gives rise to its complex functional dynamics. Here, we address this challenge by examining the inter-ictal activity of five patients with medically refractory epilepsy during ∼ 15 hours of multi-channel intracranial recording. By constructing a pairwise maximum entropy model (MEM) of the observed neural dynamics, we seek to uncover the fundamental relationship between functional activity and its underlying structural substrate. Despite only incorporating the pairwise correlations in the observed neural activity, we find that the pairwise MEM robustly fits large-scale patterns of inter-ictal power dynamics across a wide range of frequency bands, notably displaying time-invariance and cross-frequency similarity. Furthermore, across all frequency bands, we demonstrate that the pairwise MEM accurately identifies the structural white matter connections between brain regions, outperforming other common model-free measures of functional connectivity. Together, our findings show that a simple pairwise MEM, which is explicitly ignorant of higher-order correlations between three or more brain regions, not only captures complex spatiotemporal patterns of neural activity across the frequency spectrum, but also suggests that the network of structural connections in the human brain is a plausible scaffold capable of supporting observed wide-band neural dynamics.

## Introduction

An age-old question in neuroscience concerns the enigmatic relationship between the brain’s large-scale function and its underlying white matter structure (Bassett et al., 2018; Bassett and Sporns, 2017). How does the complex array of functional dynamics that we observe in the brain emerge from the static architecture of structural connections between neurons and brain regions? This question lies at the heart of several ongoing efforts, from understanding the brain’s cognitive capabilities and development, to localizing and treating human disease. Previous studies demonstrate that network models of functional connectivity (FC), calculated over several minutes of resting state fMRI, share important organizational features with network models of the underlying white-matter structural connectivity (SC) (Skudlarski et al., 2008; Hagmann et al., 2008; Honey et al., 2009; Hermundstad et al., 2013, 2014; Van den Heuvel and Sporns, 2013; Wang et al., 2015). Additionally, recent research shows that accounting for indirect and higher-order structural connections between brain regions can further improve predictions of the FC that is supported (Goñi et al., 2014; Becker et al., 2018). Coupling between FC and SC is not constant over time, but rather evolves throughout adolescence (Hagmann et al., 2010), and is notably altered in disorders of mental health (Skudlarski et al., 2010; Zhang et al., 2011). Moreover, studies have discovered hierarchical and small-world organization in both the functional connectivity between brain regions as well as in the network of structural wires connecting them (Meunier et al., 2009b, 2010; Bassett et al., 2008; Bassett and Bullmore, 2006), and the dynamics of these functional and structural networks have been implicated in cognition (Braun et al., 2015; Bassett et al., 2011; Shine et al., 2016; Bassett et al., 2015), development (Supekar et al., 2009; Dosenbach et al., 2010; Meunier et al., 2009a), and psychiatric conditions (Micheloyannis et al., 2006; Menon, 2011; Yerys et al., 2017).

Commonly used measures of FC, such as the Pearson correlation, can reveal emergent patterns of neural activity at short time scales (e.g., Ashourvan et al. (2017)) and can also provide an accurate mapping of the brain’s functional organization at longer time scales (Kramer et al., 2011, 2010). Such model-free measures of FC, however, fundamentally quantify the functional similarity between neural units (e.g., neurons or brain regions), and therefore are limited in their ability to discern between different underlying mechanisms (Adachi et al., 2011; Lu et al., 2011). For example, given two brain regions with a high Pearson correlation, one cannot distinguish between the following three scenarios: (i) The regions are communicating directly via a structural connection, (ii) the regions are communicating indirectly via a structural pathway bridging intermediate regions, or (iii) the two regions are not communicating at all, but are rather being driven by a common third region. To distinguish between direct and indirect communication, and between pairwise versus higher-order interactions, one must begin with a model of how patterns of activity are generated in the brain, and then infer the network of underlying interactions that best describes correlations in the data.

One model of neural activity – the pairwise maximum entropy model (MEM) – has generated particular interest, primarily because it is formally minimal in the sense that it accounts for the observed pairwise correlations in the data while remaining explicitly ignorant with regard to all higher-order correlations (Jaynes, 1957). In recent studies, this model has proven sensitive to the spatiotemporal patterns of co-activation in neuronal spiking at the micro-scale (Schneidman et al., 2006; Shlens et al., 2006; Yeh et al., 2010) and in patterns of blood-oxygen-level-dependent (BOLD) activity at the macro-scale (Roudi et al., 2009; Watanabe et al., 2013, 2014; Ashourvan et al., 2017). The pairwise MEM is based on the principle of maximum entropy, which states that the probability distribution (i.e., the model) that best represents one’s current state of knowledge about a system is the one with the largest *entropy* (or *uncertainty*) in the context of previously observed data. Fitting a pairwise MEM entails iteratively adjusting the individual activation of regions as well as the interaction strength of all region pairs until the estimated covariance matrix matches the covariance observed in the data. In this way, the MEM makes quantitative predictions about the frequencies of global activity patterns, rather than simply quantifying the similarities between regions, as is common in studies of FC. A good fit of the pairwise MEM indicates that the observed patterns of power amplitude states can be accurately described with a combination of first-order activations and second-order (pairwise) co-activations between electrodes. A poor fit signifies that higher-order effects (e.g. interactions between triplets or quadruplets of regions) and/or common inputs from other external regions may be required to explain the behavior of the system. Notably, prior studies using resting state fMRI data have demonstrated that the pairwise MEM accurately predicts the observed patterns of regional activations, and provides a more accurate map of the underlying structural connectivity (derived from diffusion tensor imaging (DTI)) than conventional FC-based methods (Watanabe et al., 2013).

Here, we build on prior work by examining intracranial EEG (iEEG) power dynamics across several frequency bands, ranging between 4–180 Hz, using a pairwise MEM as our measure of FC. We consider iEEG data acquired from five patients with medically refractory epilepsy over 14.6 ± 2.5 hours (in one-hour segments) of inter-ictal recording. Unlike scalp EEG, which suffers from noise and selective high frequency filtering (Grech et al., 2008), iEEG provides a unique opportunity to record relatively focal neural activity with high temporal resolution from distributed brain structures across a wide range of frequencies. We define ‘on’-’off’ activation states for each brain region by binarizing the normalized envelope of the power amplitude of each frequency band (Figure 1). Our use of the power amplitude envelope to define the activation states is motivated by several factors. First, it has been shown that the BOLD fMRI signal echoes the envelope of high frequency neural activity (Logothetis et al., 2001; Winder et al., 2017), specifically when measured by iEEG (Lachaux et al., 2007; Jacques et al., 2016; Nir et al., 2008; Ojemann et al., 2013; Betzel et al., 2018; Reddy et al., 2018). Thus, in light of studies linking BOLD fMRI FC to white matter SC (Honey et al., 2009; Watanabe et al., 2013), we hypothesize that the power of iEEG recordings should also exhibit a clear relationship with the underlying SC. Second, a large body of evidence has demonstrated that iEEG oscillations play an important role in healthy cognitive function, and that breakdowns in oscillatory power are linked to cognitive disorders (Fries, 2009; Wang, 2010; Uhlhaas and Singer, 2006). Third and finally, by defining our activity states using band-passed power amplitudes, we gain the ability to analyze patterns of activity across both time and frequency, thereby avoiding inconsistent phase relationships between electrodes caused by spatially non-stationary signal sources, such as iEEG spiral waves (Huang et al., 2010).

**Figure 1.**
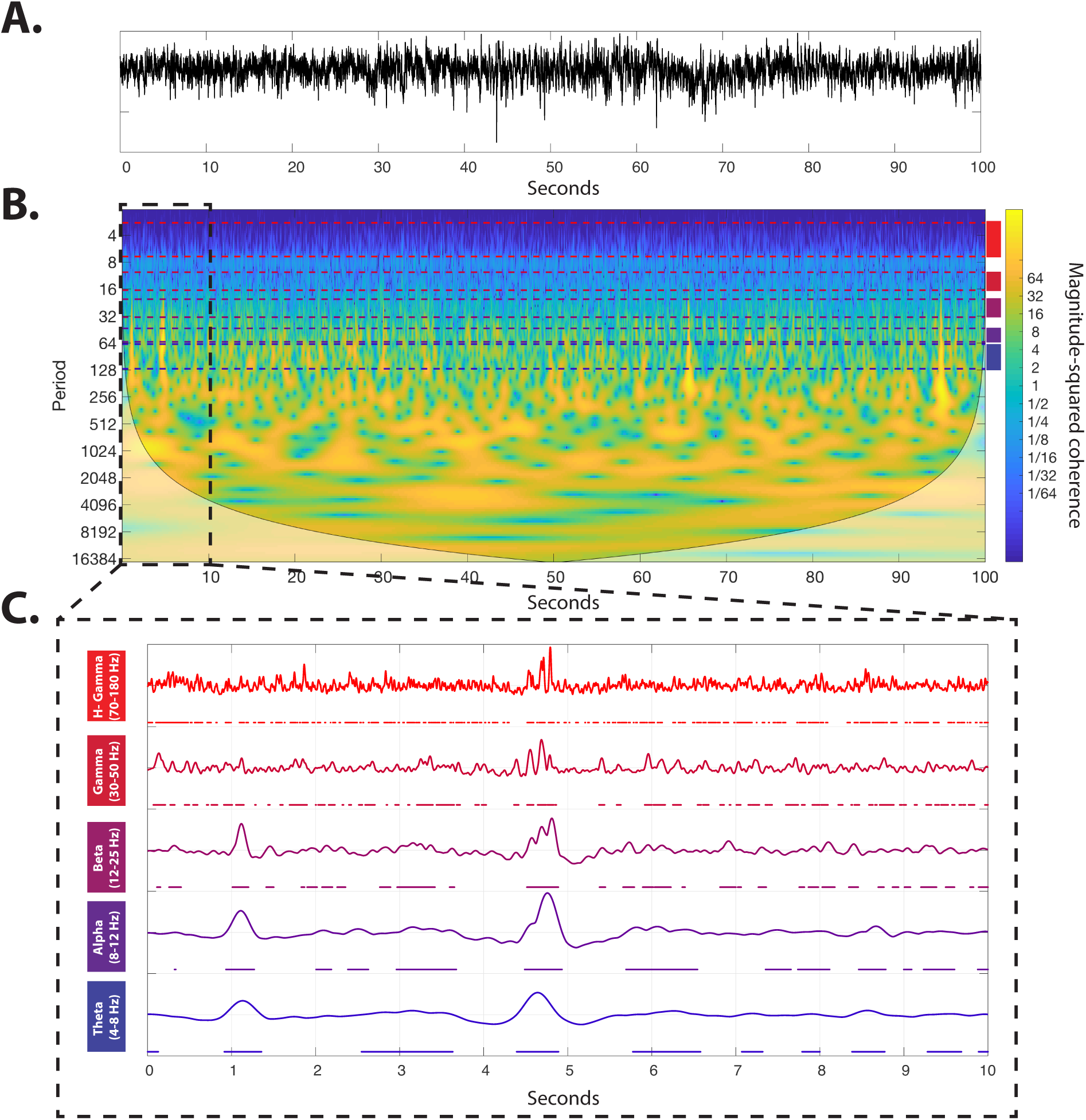
Construction of power amplitude states from iEEG time-series. ***(A)*** The raw iEEG time-series recorded from a sample electrode. ***(B)*** A wavelet decomposition of 100 seconds of raw iEEG time-series (Grinsted et al., 2004). Color-coded dashed lines show the *θ, α, β, γ*, and high *γ* frequency bands. ***(C)*** The average power of all frequencies within each band is high-pass filtered at 0.5 Hz, and normalized to obtain a *z*-score. The resultant time series were then thresholded (*>*0) separately for each band, to create the binarized power amplitude states on which the MEM operates. Note that the ‘on’ states are marked by color-coded dots under each curve.

Our results reveal that across all patients, the estimated interaction matrices from the pairwise MEM are more consistent across both time and frequency than the corresponding FC matrices defined simply by pairwise correlations. These results suggest that the intrinsic interactions between brain regions may remain relatively stationary over time, and that a common underlying mechanism may support functional dynamics across all frequency bands. We hypothesize that the physical wiring of SC likely comprises this common scaffold, and we test this hypothesis by comparing the network topology of the inferred maximum entropy interactions with the architecture of structural connections between brain regions. Receiver operator curves (ROCs) reveal that the pairwise MEM robustly identifies the presence or absence of structural connections between regions across all frequency bands for all patients. Furthermore, although the FC defined by power amplitude correlations in the *θ* and *α* bands allow identification of the underlying SC in a few patients, the pairwise MEM generates a consistently stronger prediction of SC across all frequency bands and all patients. Despite the similarity between the empirically observed covariance structure and that predicted by the pairwise MEM, we find that empirical observations of large-scale iEEG states often diverge from MEM predictions, notably in *γ* bands. These observations suggest that higher-order interactions and/or external sources may contribute to the observed activation states. Despite this divergence in the *γ* bands, our findings generally suggest that regional activation and pairwise co-activation, supported by an underlying scaffold of structural wiring, can explain a large portion of the power amplitude patterns across a wide range of frequencies (4–180 Hz). These results offer new avenues for clinical interventions based on targeted alteration of a patient’s structural connectome, with the ultimate goal of influencing neurophysiological dynamics while preserving the cortical real estate and minimizing damage to SC as documented in more aggressive surgical options (such as callosotomy (Concha et al., 2006; Song et al., 2003; Concha et al., 2007; Yogarajah et al., 2010; Taylor et al., 2018)).

## Results

### The pairwise MEM exhibits high cross-frequency similarity and temporal stationarity

Functional connectivity representing correlations in iEEG states has been shown to display patterns that vary significantly across both frequency and time (Chu et al., 2012; Kramer et al., 2010, 2011). By contrast, given our hypothesis that the inferred maximum entropy parameters should reflect the underlying SC, one would expect these pairwise interactions to maintain a similar architecture across frequency bands. Indeed, the matrix *J*_*i j*_, which encodes the inferred pairwise interactions between electrodes, displays greater cross-frequency similarity than the corresponding FC matrices representing Pearson correlations in power amplitude. Specifically, we found that 4 out of the 5 subjects displayed significantly higher average pairwise correlation between *J*_*i j*_ matrices across frequency bands than that of power amplitude FC matrices (*t*-test, *p* = 6.30 × 10^−3^, *p* = 6.53 × 10^−4^, *p* = 0.78, *p* = 1.39 × 10^−6^, *p* = 2.54 × 10^−9^, respectively; SI-Figure 1). Furthermore, the average interaction strengths in the *J*_*i j*_ matrices did not differ significantly across most frequency bands (*t*-test, FDR p*<*0.05); the notable exception is the high *γ* band, wherein the average interaction strength was significantly larger than in other frequencies bands. Interestingly, the relation between the estimated pairwise interactions and the Euclidean distances between electrodes was consistent across frequency bands, while an analogous relation between power amplitude FC and Euclidean distance was only observed in the low frequency band (see SI-Figure 2). Consistent with prior studies demonstrating that iEEG FC is relatively stable when estimated over large time windows (Chu et al., 2012), we found that both the pairwise maximum entropy interactions and the power amplitude FC, when estimated over 1 hour intervals, were consistent across time. However, a statistical comparison between the pairwise correlations of FC matrices (over all possible one-hour segments) and that of *J*_*i j*_ matrices, revealed that the pairwise MEM interactions were significantly more stable across all patients and frequency bands (*t*-test, p *<*0.05; SI-Figure 3). Together, these results support the notion that brain regions’ intrinsic propensity for interactions are similar across a wide range of frequencies and display time-invariance over several hours.

### The pairwise MEM estimated from iEEG predicts structural connectivity estimated from diffusion imaging

In contrast to common FC measured based on Pearson correlations, the pairwise MEM attempts to explain patterns of activity in the data by identifying underlying interactions between pairs of regions that can account for the observed correlations. Because the pairwise MEM takes into account the indirect propagation of influence between the electrodes, we hypothesized that the estimated maximum entropy interaction matrices would be sparser than the FC matrices obtained from power amplitude correlations. Moreover, we hypothesized that the pairwise MEM would identify underlying white-matter tracts that support the observed functional activity. Consistent with our first hypothesis, we find that the *J*_*i j*_ matrices are notably sparser than the FC matrices obtained from the power amplitude correlations, and display a distribution of strengths that is more skewed (see SI-Figure 2). Additionally, consistent with our second hypothesis, we find that the *J*_*i j*_ matrices are highly predictive of the underlying (binarized) SC across all frequency bands, as determined by a receiver-operator curve (ROC) analysis (see Figure 2.A–D for results obtained in a single representative subject, and see Figure 3 for consistent results across all subjects). The ability of the pairwise MEM to identify the underlying SC is even more pronounced when noting that the FC matrices, obtained from both power amplitude correlations and traditional time series correlations, are significantly less predictive of white matter SC (*t*-test, FDR p *<* 0.05; SI-Figure 4) across all frequencies, patients, and SC binarization thresholds (SI-Figure 5). Furthermore, across all frequency bands, 4 out of the 5 subjects displayed a significant positive relationship between the weights of the SC matrices and the weights of the interaction matrices obtained from the pairwise MEM (SI-Figure 6). Together, these results provide converging evidence that the pairwise MEM robustly uncovers the underlying structural connectivity between regions and, conversely, that this structural wiring provides a scaffold that supports the observed wide-band cortical dynamics.

**Figure 2.**
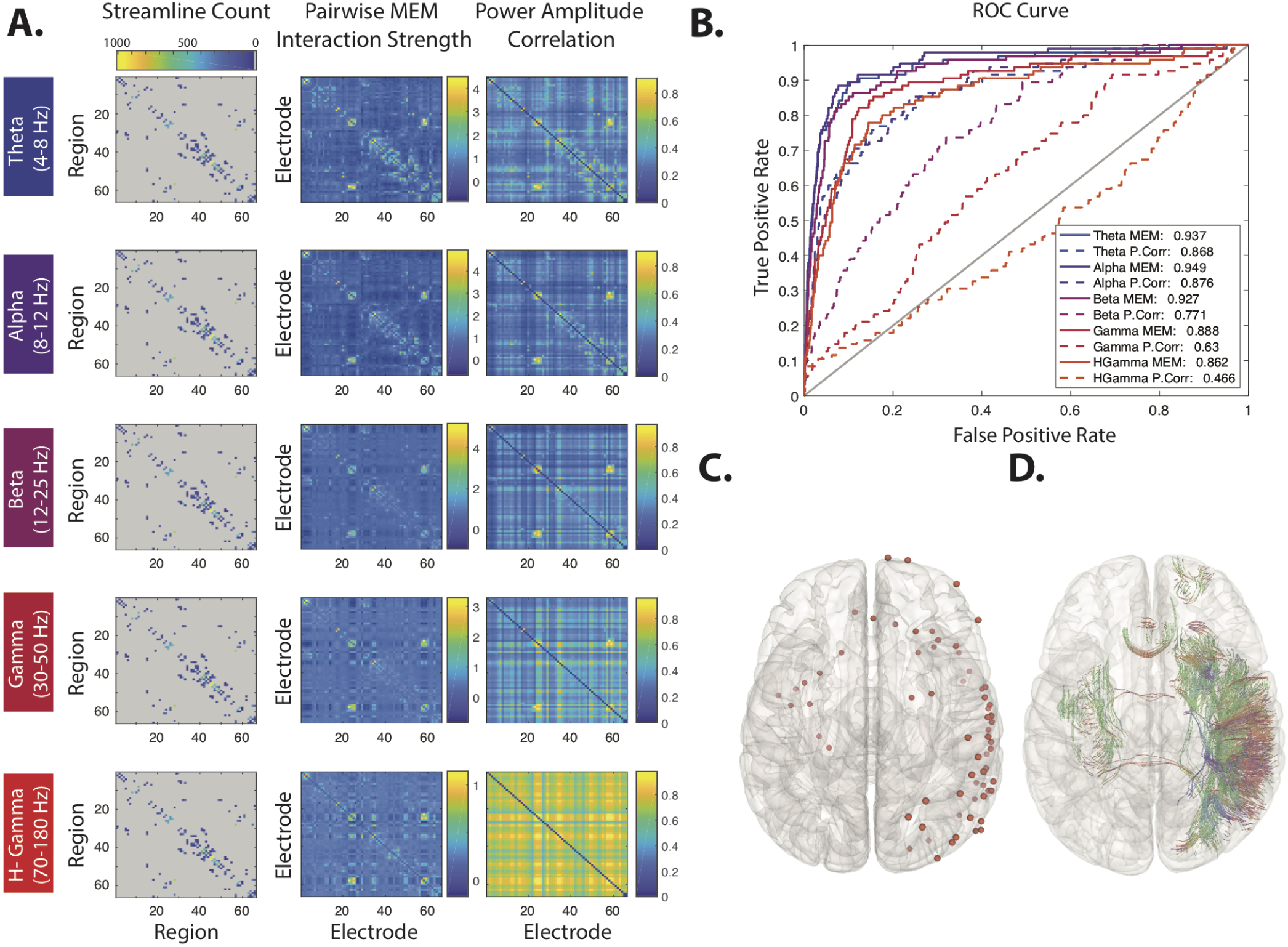
Predicting structural connectivity from estimates of functional connectivity. ***(A)*** *(Left Column)* A structural connectivity matrix from a single representative patient (#5), each element of which indicates the number of streamlines estimated between a pair of brain regions. *(Middle Column)* The interaction matrices, *J*_*i j*_, obtained from the pairwise MEM. *(Right Column)* The functional connectivity matrices estimated from band-passed power amplitude correlations. For regions covered by multiple electrodes, we selected only the electrode closest to the region’s centroid for display in these plots as well as in the ROC analysis. ***(B)*** ROC curves for identification of anatomically connected regions based on interaction matrices (solid lines) and band-passed power amplitude correlations (dashed lines). Lines are color-coded based on the frequency band. ***(C)*** Brain overlay providing the position of the selected electrodes (red dots). ***(D)*** Streamlines connecting the regions covered by the selected electrodes in the same patient. Here the 3-dimensional orientations of the streamlines are coded by color.

**Figure 3.**
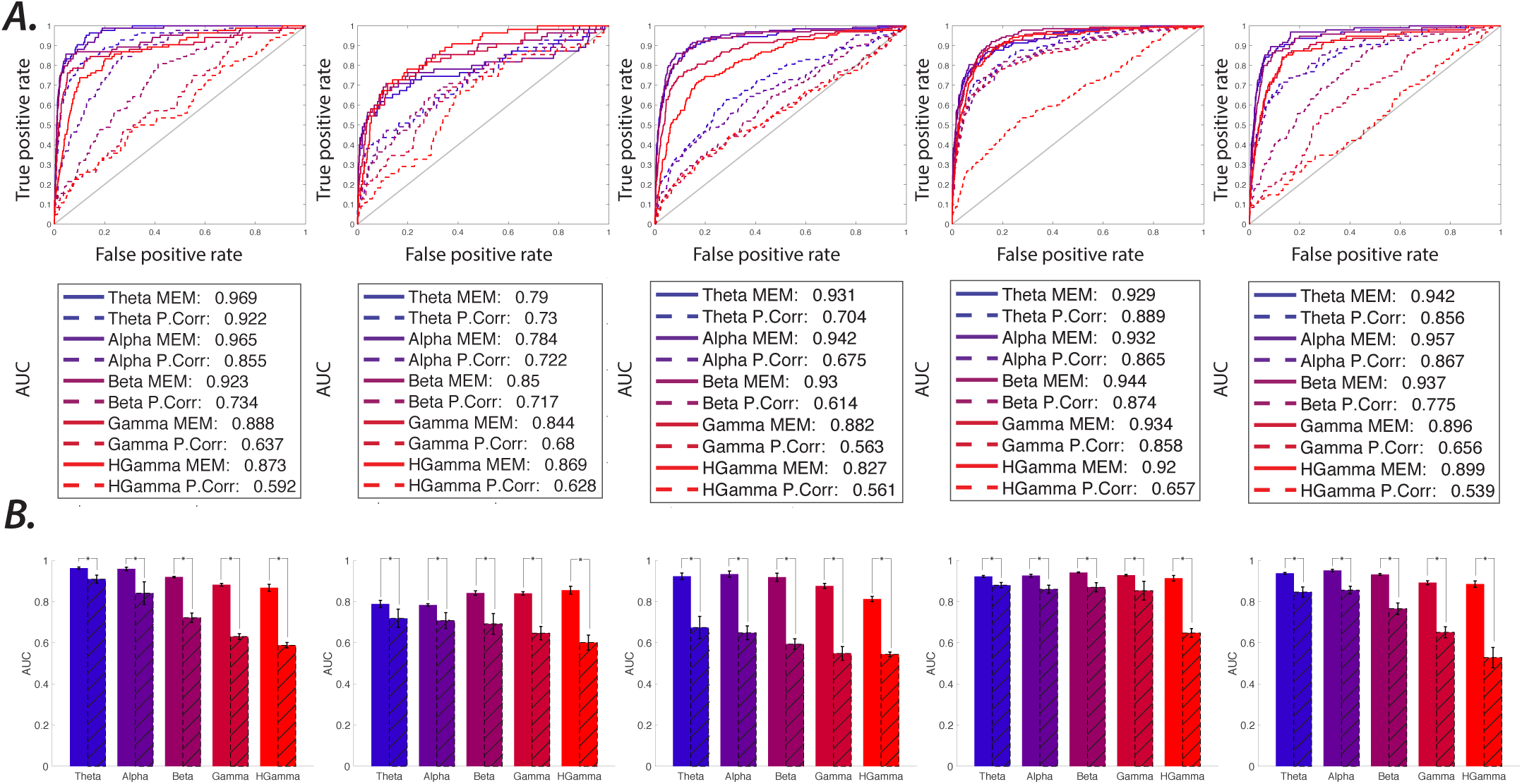
ROC analysis for the detection of SC based on functional interaction estimates. ***(A)*** ROC curves for the identification of anatomically connected versus unconnected ROI pairs (i.e., binarized SC) based on the average (14.6 ±2.5 hours of one-hour segment data per patient) *J*_*i j*_ interaction matrices (solid lines) and average power FC (dashed lines) for all 5 patients (columns). Lines are color-coded based on frequency band. The bottom tables provide the AUC values for all curves. ***(B)*** The color-coded bars represent the mean AUC and error bars represent the standard deviation (across all one-hour segments) calculated for the SC matrices for different frequency bands. The average AUC calculated using *J*_*i j*_ interaction matrices (sold bars, left) are significantly (*t*–test, p *<* 0.05 FDR, marked by ‘*’) higher than that calculated using power FC matrices (dashed bars, right) across all frequency bands. Similar results are found at even higher SC pruning thresholds, as shown in SI-Figure 3.

### Accounting for distance in the relation between the pairwise MEM and structural connectivity

Extensive prior work has demonstrated that the weights of structural connections between brain regions decreases as a function of inter-regional distance (Donahue et al., 2016; Kaiser and Hilgetag, 2004; Lewis et al., 2008; Rubinov et al., 2015). The effects of distance on SC are even more pronounced with respect to iEEG data because the electrode grids and strips are distributed focally rather than uniformly across the brain. Indeed, we observe that the most structurally connected electrode pairs are proximal and that long range fibers are scarce (SI-Figure 7.A). In fact, the distance between electrodes is highly predictive of the presence of a structural connection between them (average AUC across subject, = 0.986 ± 0.003). In order to better account for the role of distance, we repeated the ROC analysis after removing a variable percentage of the closest electrode pairs (SI-Figure 7.B). With the exception of one subject’s data in the *θ* and *α* bands, we observed that the interaction matrices estimated by the pairwise MEM continued to provide more accurate predictions of SC than the FC matrices obtained from power amplitude correlations. In a complementary approach, we regressed out the inter-regional Euclidean distances from both the maximum entropy interaction strengths as well as the power amplitude correlations. We then repeated the ROC analysis, finding, across all frequency bands, that the FC matrices obtained from power amplitude correlations did not significantly predict the presence of SC (SI-Figure 7.C). By contrast, this distance effect is markedly weaker for the interaction matrices obtained from the pairwise MEM, with the exception of the high-*γ* band in 4 out of the 5 subjects. Together, these results suggest that, although the presence of SC is highly confounded with inter-regional distance, especially in the high-*γ* band, the pairwise MEM is able to detect direct structural connections between brain regions beyond those anticipated by inter-regional distance alone.

### The pairwise MEM robustly predicts the observed covariance structure of power amplitude states

Because a large portion of the cortex is canvassed by electrodes in our patient sample, we have the necessary coverage to measure interactions between distant brain regions. However, we expect to see some degree of divergence between the observed and predicted state probabilities for three reasons: (i) increasing the dimensionality of the system in general decreases the accuracy of a fit due to an exponential increase in the state space, (ii) unmeasured brain regions cannot be included in the model and are therefore likely to induce systematic errors, and (iii) real-world neural activity is likely to exhibit some degree of higher-order interactions. We assessed the performance of the pairwise MEM first by comparing the empirically observed covariance matrices with the computationally estimated covariance matrices obtained from the model. We estimated the model-based covariance matrices by randomly sampling the probability distribution using the Metropolis-Hastings algorithm (see Methods). We found that the estimated covariance matrices are statistically similar to the empirical covariance matrices across all frequency bands (average correlation values between empirical and estimated covariance matrices for all patients, 0.96±0.01, 0.95±0.07, 0.94±0.03, 0.98±0.01, 0.96±0.04, *p <* 0.001 Bonferroni correction, respectively). Differences between the estimated and empirical covariance matrices existed along the diagonal, where estimated values are significantly larger than empirically measured values (all patients and frequency bands, paired *t*–test, *p <* 0.001 Bonferroni correction, see SI-Figure 8), and also among pairs of electrodes located near one another physically (as seen in Figure 4). We also observed a rapid decrease in estimation errors as a function of the distance between electrodes across all frequency bands (see Figure 4.E for results obtained from a single subject, and see SI-Figure 9 for results obtained from all subjects). The similar and highly significant (*p* ≈ 0) exponential fit to the covariance estimation error versus electrode pair distance across all subjects and frequency bands highlights the much higher estimation error for nearby electrode pairs (SI-Figure 9). In summary, these results demonstrate that the pairwise MEM provides a statistically robust prediction of the observed covariance structure of the power amplitude states across all frequency bands, and that its relatively small approximation errors are predominantly local.

**Figure 4.**
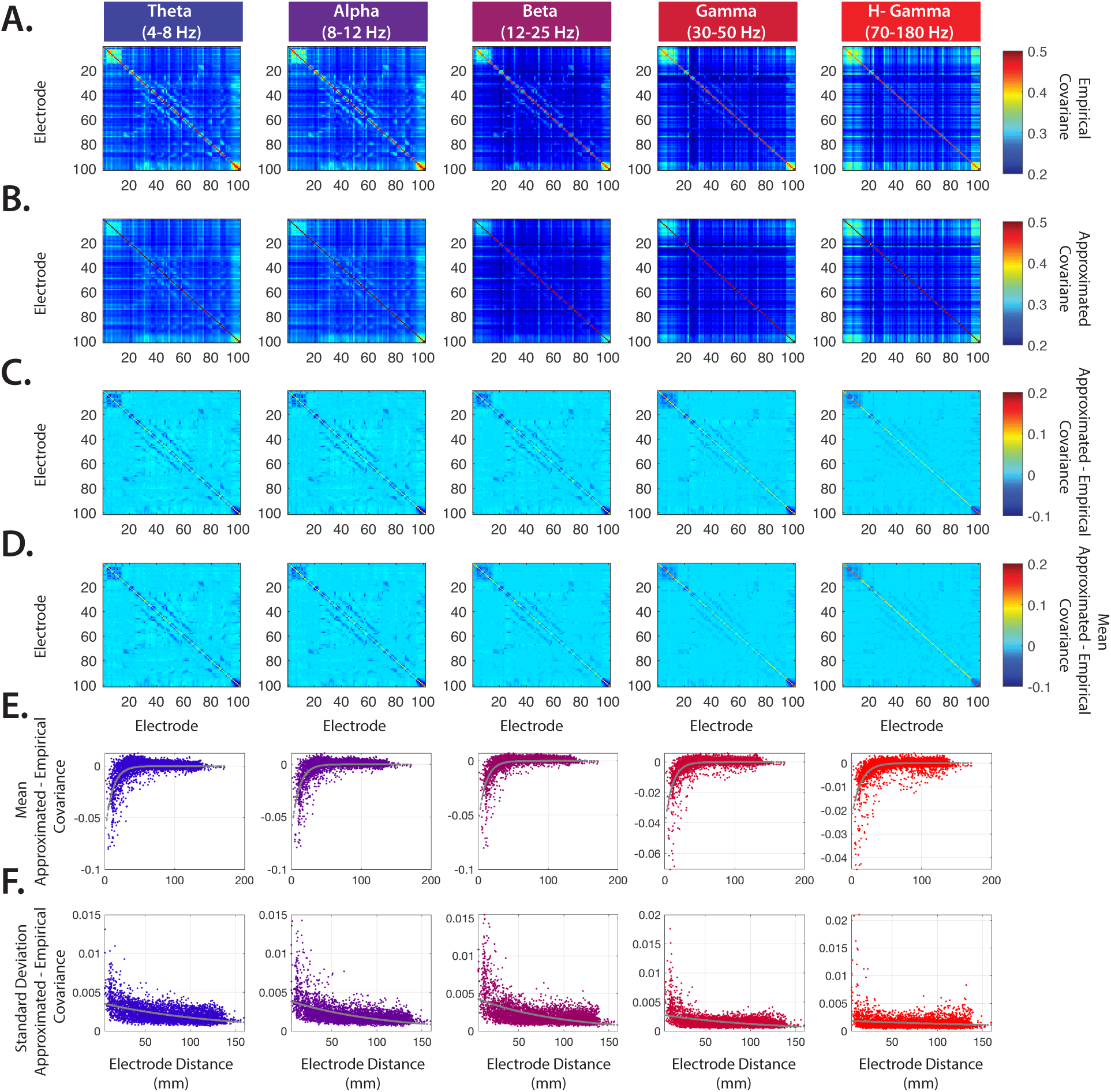
Comparison of covariance matrices of neural activity with those predicted by the pairwise MEM. ***(A)*** The covariance matrices calculated from band-passed power amplitude states from a single representative patient (#5). Each column contains results obtained from a distinct frequency band. ***(B)*** The covariance matrices estimated from the pairwise MEM. ***(C)*** Differences between the empirically observed covariance matrices and the computationally estimated covariance matrices obtained from the pairwise MEM for a single representative patient (#5). ***(D)*** The average difference matrices across all 19 data sets, reflecting the approximation error. ***(E)*** The average approximation error as a function of the pairwise Euclidean distance between electrodes, fit by an exponential function (green dashed line). Note that the error is notably higher between nearby electrodes. For convergent results across all subjects, see SI-Figure 9. ***(F)*** The standard deviation of the approximation error as a function of the pairwise Euclidean distance between electrodes.

Another way to assess the goodness of fit for the pairwise MEM is to calculate the degree to which the estimated probabilities for the observed activity states diverge from the empirically measured probabilities. As a baseline, this divergence between the pairwise model and the observed statistics is often compared to the divergence of the simpler first-order model, which assumes that each brain region is behaving independently from the others. It is important to note that this calculation can be computationally expensive for large systems because the state space increases exponentially (= 2^*n*^) with the number of electrodes (*n*). To circumvent this issue, we instead compared the probability of all empirically observed states to their predicted probabilities. SI-Figure 10 demonstrates the degree of divergence between the empirical and estimated probabilities of all empirically observed states for a sample one-hour segment across all frequency bands. As expected, the largest estimation errors are observed in states with the lowest empirical probabilities. However the model produces notably large errors for a large number of states with relatively high empirical probabilities, specifically in the *γ* and high-*γ* bands (SI-Figure 10.A). Additionally, we examined the goodness of fit of the model for higher binarization thresholds of the power amplitude time series (SI-Figure 10.A). These results show that although higher thresholds provide different distributions of approximation errors, the model still provides a relatively poor prediction for a large number of states, consistent with the presence of higher-order interactions. It will be interesting in future to assess the presence and extent of such higher-order interaction in the *γ* and high-*γ* bands using higher-order MEMs Ganmor et al. (2011). Nevertheless, the ability of the estimated interaction matrices to predict the underlying SC is highly robust to all binarization thresholds, despite the difference in the patterns of approximation errors (SI-Figure 10.B-C and SI-Figure 11). Together, these results suggest that a simple model using only a combination of regional activation rates and pairwise co-activation rates can closely approximate the observed covariance structure in the iEEG power time-series.

## Discussion

Here, we constructed a series of pairwise MEMs capable of reproducing the covariance structure observed in iEEG power amplitude states recorded from 5 patients with partial-onset refractory epilepsy. Careful examination of the covariance matrices predicted by the pairwise MEMs revealed markedly small approximation errors across all frequency bands. Nevertheless, we identified divergence between the estimated and empirical probabilities of states, mostly in the *γ* bands, suggesting that *γ* oscillations may be strongly driven by higher-order interactions. These results suggest that, while external common sources and higher-order interactions may contribute to the observed neural activity, the activation and pairwise co-activation rates between brain regions capture most of the observed variance. Importantly, our results also revealed that the inferred pairwise interactions in the MEM were highly stable over time and maintained a high degree of similarity across frequency bands. Notably, the estimated interaction matrices were both more consistent over time and more similar across frequencies than a commonly used measure of FC, the Pearson correlation between the power amplitude time series. We then investigated the relationship between the pairwise maximum entropy interactions, which were calculated only to account for the functional iEEG data, and the underlying white matter connectivity between regions by performing ROC analysis. AUC results revealed that the pairwise MEM provides accurate predictions of the underlying structural connectivity across all subjects and frequency bands, an effect similar to – though significantly stronger than – that observed in power amplitude FC at low frequencies. Taken together, these findings demonstrate the utility of the pairwise MEM for explaining covariance patterns in iEEG signals and in uncovering their underlying anatomical substrates.

### The structural connectivity scaffold for functional connectivity

A critical open question in neuroscience is how the relatively fixed white matter structure of the brain gives rise to complex functional dynamics. Some have argued that these complex dynamics are well-approximated by a single underlying process (Laumann et al., 2016; Liegeois et al., 2017), while others favor a model comprised of multiple processes that the brain can switch between (Friston et al., 2003; Deco et al., 2013; Laumann et al., 2016). In either scenario, it is critical to understand from a statistical perspective whether the observed dynamics can be explained by a fixed structural scaffold, such as that represented by the pattern of white matter fiber bundles connecting large-scale brain regions. The answer depends to some degree on whether or not one can identify statistical features of the functional dynamics that display relatively time-invariant properties. Some evidence for such time-invariance has recently been obtained from eletrocorticography, where patterns of functional connectivity between electrodes appear relatively stable in windows larger than a few minutes (Chu et al., 2012; Kramer et al., 2010, 2011). These data are consistent with our observations that patterns of interactions estimated over one-hour windows were relatively stable across several hours. Notably, this temporal stability was markedly higher in the interactions estimated from the pairwise MEM than in the functional connections estimated from the Pearson correlation in power amplitude time series. Our results therefore provide evidence of high stationarity of the intrinsic pairwise relationships between brain regions that gives rise to power amplitude dynamics, and further motivate a careful investigation into the correspondence between those relationships and the underlying anatomical projections.

A second line of evidence supporting the existence of a common structural scaffold for the observed dynamics is the presence of notable similarity in the estimated pairwise interactions across frequency bands. This statistical similarity is reminiscent of the cross-frequency interactions that have been observed in other studies, and that are thought to play a role in integrating information across brain structures that are functionally specialized and spatially segregated (Canolty et al., 2006; Hyafil et al., 2015; Lisman and Jensen, 2013). It is commonly thought that the diverse rhythms present in different frequency bands are associated with different spatio-temporal scales of neural activity (Raghavachari et al., 2006; Canolty et al., 2007), with low frequencies driving activity over larger spatial areas and high frequencies driving activity over smaller spatial areas (Von Stein and Sarnthein, 2000). Our work offers a useful complement to these prior studies by demonstrating that the pairwise MEM provides higher cross-frequency similarity in estimated interactions than conventional measures of functional connectivity. Future work could examine the real-time dynamics of cross-frequency coupling by combining power amplitude states across all bands in a single model. Taken together, our observations of high cross-frequency similarity indicate that a common underlying physiological (Manning et al., 2009) or anatomical (Betzel et al., 2018) mechanism is likely driving the functional interactions between brain regions across all frequency bands.

Our observation of the relatively high cross-frequency similarity and stationarity of interaction matrices estimated from the pairwise MEM is consistent with the notion that the patterns of interactions track an underlying structural scaffold. We validate this possibility with rigorous statistical testing, showing that the interactions can be used to predict underlying white matter structure in the same patients, and that these predictions are stronger for the pairwise MEM interactions than for a conventional measure of functional connectivity, consistent with prior work in other imaging modalities (Watanabe et al., 2013). The higher structure-function coupling obtained with the MEM could be due to the fact that the MEM simultaneously fits all regional activation rates and all pairwise interactions. Partial correlation approaches that operationalize the latter also tend to show high structure-function coupling (Salvador et al., 2005; Watanabe et al., 2013). It will be interesting in future work to consider the relevance of the MEM for explaining the effects of stimulation, for example where single external pulse stimulation of cortex propagates to mono-and poly-synaptically connected regions via cortico-cortical and cortico-subcortical projections (Steriade and Amzica, 1996; Matsumoto et al., 2006). In sum, our results suggest that the functional interactions facilitated by the direct underlying structural connectivity between brain regions largely shapes the complex broadband functional dynamics evident in power amplitude time series.

### Statistical accuracy of the pairwise MEM and potential biophysical mechanisms

Pairwise MEMs have been employed effectively to explain patterns of collective neural activity across a range of spatial scales (Schneidman et al., 2006; Watanabe et al., 2013; Ashourvan et al., 2017). Here, we demonstrate that the pairwise MEM allow us to approximate the empirical covariance patterns of power amplitude states in several biologically relevant frequency bands. We observed that the accuracy of the approximated covariance in the pairwise MEM was markedly high across all frequency bands. While the overall accuracy was high, we sought to better understand where and why the pairwise MEM produced its greatest errors. We observed that the model systematically overestimated electrode activation rates while underestimating electrode-to-electrode co-activation rates, particularly between electrodes that were located in close physical proximity, the latter observation being consistent with prior work (Raghavachari et al., 2006; Canolty et al., 2007; Nunez and Srinivasan, 2006; Dickson et al., 2000). Additional examination of the empirical and estimated probabilities revealed notable divergence in the probabilities of a number of the observed states, with this divergence being most pronounced in the *γ* and high *γ* bands. Although some approximation error is expected due to our heuristic algorithm for calculating pairwise MEM parameters, we speculate that the error that we observe might also originate from biophysical mechanisms contributing to the co-activation of nearby electrodes. Examples of such biophysical mechanisms include the well-known and markedly strong white matter connectivity between nearby brain regions (Betzel and Bassett, 2018), spatially distributed activation sources, and spiral waves (Huang et al., 2010). Moreover, it is important to note that even though iEEG electrodes cover a large portion of the cortical surface in these patients, we are still only able to capture a small fraction of the full system, and it is highly likely that some amount of co-activation is driven by input from external sources that are not covered by an electrode. Although the MEM could be extended to include higher-order parameters beyond the pairwise interactions considered here, the practical limitation of partial brain coverage and the previously noted biophysical mechanisms combine to hinder the interpretability of higher-order mechanisms.

## Methodological considerations

One common concern regarding the use of iEEG datasets from patients with epilepsy to study the functional organization of the human brain is their documented aberrations of structural connectivity as well as structure-function coupling (Liao et al., 2010; Besson et al., 2014; Bonilha et al., 2012; Zhang et al., 2011). Since iEEG datasets from healthy controls do not exist, we cannot directly address these concerns. Nevertheless, we draw on recent work demonstrating that the patterns of ECoG functional connectivity in patients show statistical similarities to structural connectivity estimated in healthy volunteers (Betzel et al., 2018), and that this statistical similarity is upheld and even strengthened during ictal epochs (Reddy et al., 2018; Shah et al., 2018). Future work using data from patients with other pathologies, or using source-localized MEG in healthy patients, could be helpful in further understanding the nature of the structure-function correspondence accessible to the MEM.

Preprocessing methods and parameter choices can impact estimates of structural connectivity. However, we demonstrate in SI-Figure 12 that the observed correspondence between functional interactions and structural connections were robust to reasonable variation in these choices. Although the pairwise MEM demonstrates a closer correspondence than the Pearson-based FC in predicting the SC across several different DTI processing methods, our results demonstrate that the deterministic streamline count matrix reflecting SC produces the strongest structure-function coupling. Future work could test the robustness of our findings to other important preprocessing parameters such as regional size, high-pass frequency filtering of the iEEG signals, and binarization threshold of power amplitude states.

It is of interest to determine the contribution of distance to structure-function coupling. The strength of a structural connection between two brain regions tends to be negatively correlated with the distance between them, as does the strength of the functional connection between them. One possibility is that structure-function coupling is driven by a common source that is simultaneously measured by nearby electrodes. To discount this possibility at least partially, we demonstrate that the pairwise MEM can be used to accurately identify structural connections between distant electrode pairs. This observation suggests that the functional dynamics captured by the pairwise MEM extends beyond common sources or local spreading phenomena, and that the presence of a structural connection between brain regions likely plays a crucial role in shaping the emergent functional activity across a wide range of frequencies. Nonetheless, future work should aim to validate our observations and further tease out the effect of electrode distance using datasets from patients with more uniform electrode coverage.

## Conclusion

Our observations have potentially important implications for understanding large-scale functional brain dynamics as well as our ability to modulate these dynamics via stimulation or resection. We demonstrated that defining the band-passed power of the iEEG signal as a functional brain state allowed us to uncover strong structure-function coupling in large-scale networks. Therefore our results provide converging evidence of the functional relevance of iEEG signal power. Furthermore our results demonstrate that the apparent divergence between structural and functional connectivity is minimized when the intrinsic functional relationships between brain regions are not examined as independent pairs. These findings are consistent with the notion that the pairwise MEM may be particularly sensitive to structurally-driven functional relations while conventional functional connectivity methods may be particularly sensitive to non-structurally-driven functional relations that might vary appreciably over short time intervals. Finally, the observed high degree of structure-function coupling suggests that structural connectivity is a useful proxy for time-invariant functional relationships (Betzel et al., 2018; Reddy et al., 2018). This observation could be useful in the treatment of epilepsy patients, where access to the brain is traditionally limited to recording loci but could be augmented with non-invasive measurements of structural connectivity for more informed surgical planning (Shah et al., 2018). Indeed, it is intuitively plausible that computational models built to inform the modulation of abnormal functional dynamics via stimulation (Gu et al., 2015) or resection (Khambhati et al., 2016) may be able to utilize patient specific structural connectivity in place of or to augment patient specific functional connectivity.

## Materials and Methods

### Patient information

5 patients (mean age 41.6, standard deviation 4.8; 3 female) undergoing surgical treatment for medically refractory epilepsy underwent implantation of subdural electrodes for localization of the seizure onset zone. All patients had unilateral temporal lobe epilepsy, determined by comprehensive clinical evaluation and validated by seizure free one-year outcomes following temporal lobectomy.

### Intracranial EEG Acquisition

De-identified patient data was downloaded from the online International Epilepsy Electrophysiology Portal (IEEG Portal, http://www.ieeg.org) (Wagenaar et al., 2013). iEEG signals were recorded at 512 Hz at the Hospital of the University of Pennsylvania, Philadelphia, PA. Subdural electrode (Ad Tech Medical Instruments, Racine, WI) configurations consisted of linear strip and two-dimensional grid arrays (2.3 mm diameter with 10 mm inter-contact spacing). A referential montage with reference electrode distant to the seizure onset zone was used for recording iEEG signals. Inter-ictal periods were selected to be at least 6 hours away from clinically marked seizures.

### Image Acquisition

MRI data were acquired using a 3-Tesla Siemens scanner (Siemens Magnetom Trio Tim Syngo MR B17, Germany) at the Hospital of the University of Pennsylvania. Diffusion-weighted images were acquired in a single-shot echo-planar imaging, multi-shell protocol (2.5×2.5×2.5 *mm*^3^ resolution, TR = 5216 ms, TE = 100 ms, FOV = 220 mm, MB acceleration factor = 2, flip/refocus angle = 90/180 degrees, phase encoding direction = anterior to posterior). A total of 119 volumes were acquired at *b*=0 (16 volumes), b=300 (8 volumes), *b*=700 (31 volumes), and *b*=2000 (64 volumes). T1-weighted MPRAGE images were also obtained for each subject (0.94×0.94×1.0 *mm*^3^ resolution, TR = 1810 ms, TE = 3.51 ms, FOV = 240 mm, flip angle = 9 degrees, phase encoding direction = right to left).

### Structural connectivity

T1-weighted MPRAGE images were used to co-register MNI-space atlases to subject structural space via AntsRegistrationSyN (Avants et al., 2008). Similarly, we used the first b0 image from each patient’s diffusion sequence to calculate co-registration transforms from subject-specific diffusion to structural space. Using this set of transforms, we brought individualized AAL 600 atlases (Hermundstad et al., 2013) for all five patients into diffusion space for tractography. Atlas to diffusion co-registrations were performed using ANTs and FSLs FLIRT. Eddy current and motion corrections were performed using FSLs 5.0.9 EDDY patch. To mitigate susceptibility distortions, we used the subjects MPRAGE T1 structural scan. This image was first brought to diffusion space by using the inverse of our FLIRT transformation for each subject, and then contrast-inverted and intensity-matched to the DWI image using FSLMATHS. Finally, the DWI image was non-linearly transformed to the shape of the MPRAGE scan, as these acquisitions are not subject to the same susceptibility distortions observed in DWIs.

We evaluated several distinct methodological approaches, and we also assessed the consistency of our findings across them. We performed deterministic and probabilistic tractography in Camino (Cook et al., 2006) on distortion-corrected DWIs to construct adjacency matrices for structural connectivity. Deterministic tractography parameters were stringent so we could err on the side of caution in evaluating the physiological feasibility of our reconstructed white matter connections. These parameters included removing estimated fibers in areas of the brain where 90% or greater of diffusion was estimated to be isotropic versus restricted (Zhan et al., 2015), where fractional anisotropy was determined to be below 0.05, or where a T1-derived brain mask ended. We also removed fibers that curved more than 50 degrees over 5 mm to again restrict fiber reconstructions to physiological feasibility. Finally, we removed fibers shorter than 10 mm to minimize spurious short-range connections based on only a few voxels, and we also removed fibers over 100 mm to prevent looping artifactual fibers from impacting our results. Structural adjacency matrices were constructed from the number of streamlines that begin and end in each pre-defined ROI. We employed equivalent parameters for probabilistic tractography and utilized the PICO algorithm (Parker et al., 2003). Processing power limited us to seeding 400 times per voxel. Within deterministic and probabilistic tractography, we evaluated streamline count (SC), fractional anisotropy (FA), neurite density (ND), and orientation dispersion (OD) weighted adjacency matrices.

### Pairwise maximum entropy model

To form unbiased predictions for the probabilities of various functional brain states, we fit a pairwise maximum entropy model, which is motivated by the principle of maximum entropy. The principle of maximum entropy states that when estimating a probability distribution given some desired constraints, one ought to consider the distribution that maximizes the uncertainty (i.e., entropy); choosing any other distribution that lowers the entropy would assume additional information beyond our desired constraints. Fitting the pairwise MEM entails tuning the first and second-order interaction parameters between regions so that the predicted activation rates and co-activation rates match the empirically observed values. An accurate fit of the pairwise MEM implies that the observed patterns of collective activity can be understood as emerging from each region’s independent activation rate combined with regions’ joint activation rates. In other words, the pairwise MEM allows us to establish a model of iEEG functional power dynamics as a probabilistic process shaped by underlying pairwise relationships between brain regions.

We fit the pairwise MEM to the thresholded and binarized normalized power amplitude states of electrodes during one hour intervals of recording per subject (Watanabe et al., 2013). To calculate the power amplitude states, we first segmented the time-series into 36, 100-second windows (SI-Figure 1.A). Next, for each segment and each electrode we calculated the wavelet power (Grinsted et al., 2004) of the time-series (Figure 1.B). We band-passed the wavelet power by averaging the power amplitude across all the frequencies within each band. Next, the power amplitude time-series (all bands) were high-pass filtered at 0.5 Hz to minimize the effect of ultra slow fluctuations of power on the binarized states. The high-passed time series were then *z*-scored and binarized by thresholding at zero to create band-passed power amplitude states (Figure 1.C).

At each time *t*, the power amplitude state is defined by 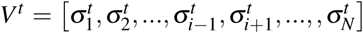 where 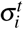 is the binarized power amplitude of electrode *i* at time *t*, where 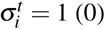 for power above (below) the threshold, and *N* is the total number of electrodes. For electrode *i*, the empirical power amplitude activation rate 〈*σ*_*i*_〉 is given by 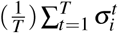, where *T* is the number of time slices. Likewise, the empirical co-activation rate between electrodes *i* and *j*, 〈*σ*_*i*_*σ*_*j*_〉, is defined by 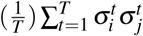.

Here, our only constraints were that the model averages 〈*σ*_*i*_〉_*m*_ and 〈*σ*_*i*_ *σ*_*j*_〉_*m*_ matched the empirical values of 〈*σ*_*i*_〉 and 〈*σ*_*i*_ *σ*_*j*_〉, respectively. Given these constraints, it is known that the probability distribution that maximizes the entropy is the Boltzmann distribution (Jaynes, 1957):

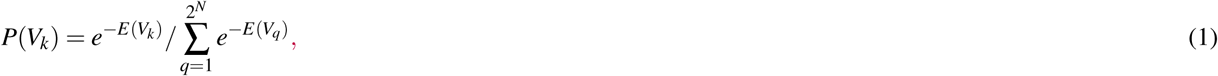

where *P*(*V*_*k*_) is the probability distribution of the *k-th* state *V*_*k*_, and *E*(*V*_*k*_) is the energy of this state, which is given by:

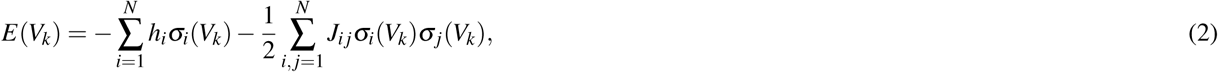

where *σ*_*i*_(*V*_*k*_) is the value of *σ*_*i*_ for state *V*_*k*_, *h*_*i*_ represents the expected base power amplitude rate of electrode *i* in isolation, and *J*_*i j*_ represents the functional interaction between electrodes *i* and *j*.

Fitting the pairwise MEM entails iterative adjustment of the parameters *h*_*i*_ and *J*_*i j*_ with a gradient descent algorithm (Watanabe et al., 2014) until the empirical averages 〈*σ*_*i*_〉 and 〈*σ*_*i*_*σ*_*j*_〉 match those in the model, 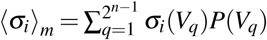 and 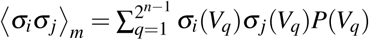. Since the state space is prohibitively large in our data, we followed (Gu et al., 2018) by approximating the model averages 〈*σ*_*i*_〉_*m*_ and 〈*σ*_*i*_*σ*_*j*_〉_*m*_ by calculating the covariance of a sequence of random samples (*N* = 10300000, after discarding the first 300000 and downsampling by a factor of 500) from the probability distribution using the Metropolis–Hastings algorithm (Metropolis et al., 1953).

### Comparison between structural connectivity and functional connectivity

We conducted ROC analysis to determine the accuracy with which the strength of the functional interactions in the pairwise MEM predict the presence of structural connectivity. We constructed the ROC curves to represent the true positive (structurally connected brain regions with functional interaction weight above the threshold) and false positive (structurally disconnected brain regions with functional interaction weight above the threshold) at different detection thresholds. The area under the curve (AUC) of the ROC plot represents the accuracy of the classification, where an AUC value of 1 indicates perfect classification, and an AUC value of 0.5 indicates classification performance at the chance level.

## Acknowledgements

This work was supported by an NINDS 1R01NS099348 to Litt and Bassett (MPI). BL also acknowledges support from the Mirowski Foundation and from Neil Barbara Smit. DSB also acknowledges support from the John D. and Catherine T. MacArthur Foundation, the Alfred P. Sloan Foundation, the Army Research Office through contract number W911NF-14-1-0679, the Army Research Laboratory through contract number W911NF-10-2-0022, the National Institute of Health (2-R01-DC-009209-11, 1R01HD086888-01, R01-MH107235, R01-MH107703, R01MH109520, 1R01NS099348 and R21-M MH-106799), the Office of Naval Research, and the National Science Foundation (BCS-1441502, CAREER PHY-1554488, BCS-1631550, and CNS-1626008). The content is solely the responsibility of the authors and does not necessarily represent the official views of any of the funding agencies.

