## Supplementary Materials for "A pairwise maximum entropy model uncovers the white matter scaffold underlying emergent dynamics in intracranial EEG"

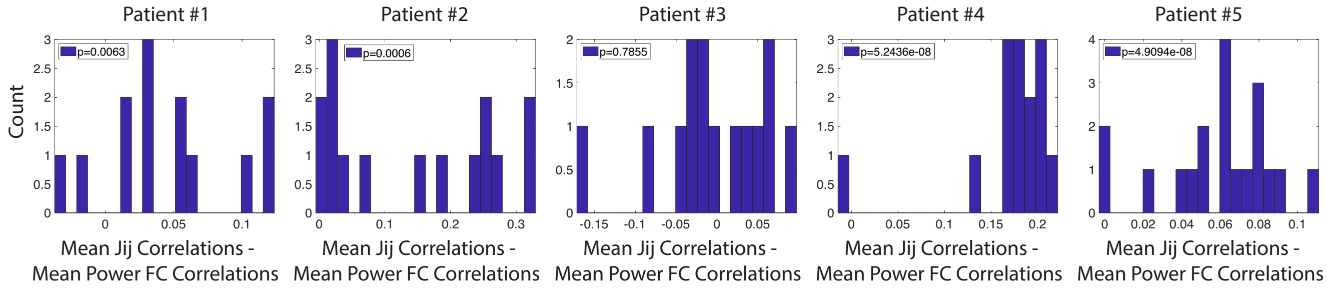


**Figure 1.** **Similarity of maximum entropy interactions across frequencies.** We calculate the correlations between the maximum entropy interactions $J_{ij}$ across all pairs of frequency bands. Each histogram shows the difference between the cross-frequency correlations in $J_{ij}$ with the cross-frequency correlations in functional connectivity. All but one patient (#3) show significantly ($t$–test, $p < 0.05$) higher average cross-frequency similarity in $J_{ij}$ matrices than in FC matrices.

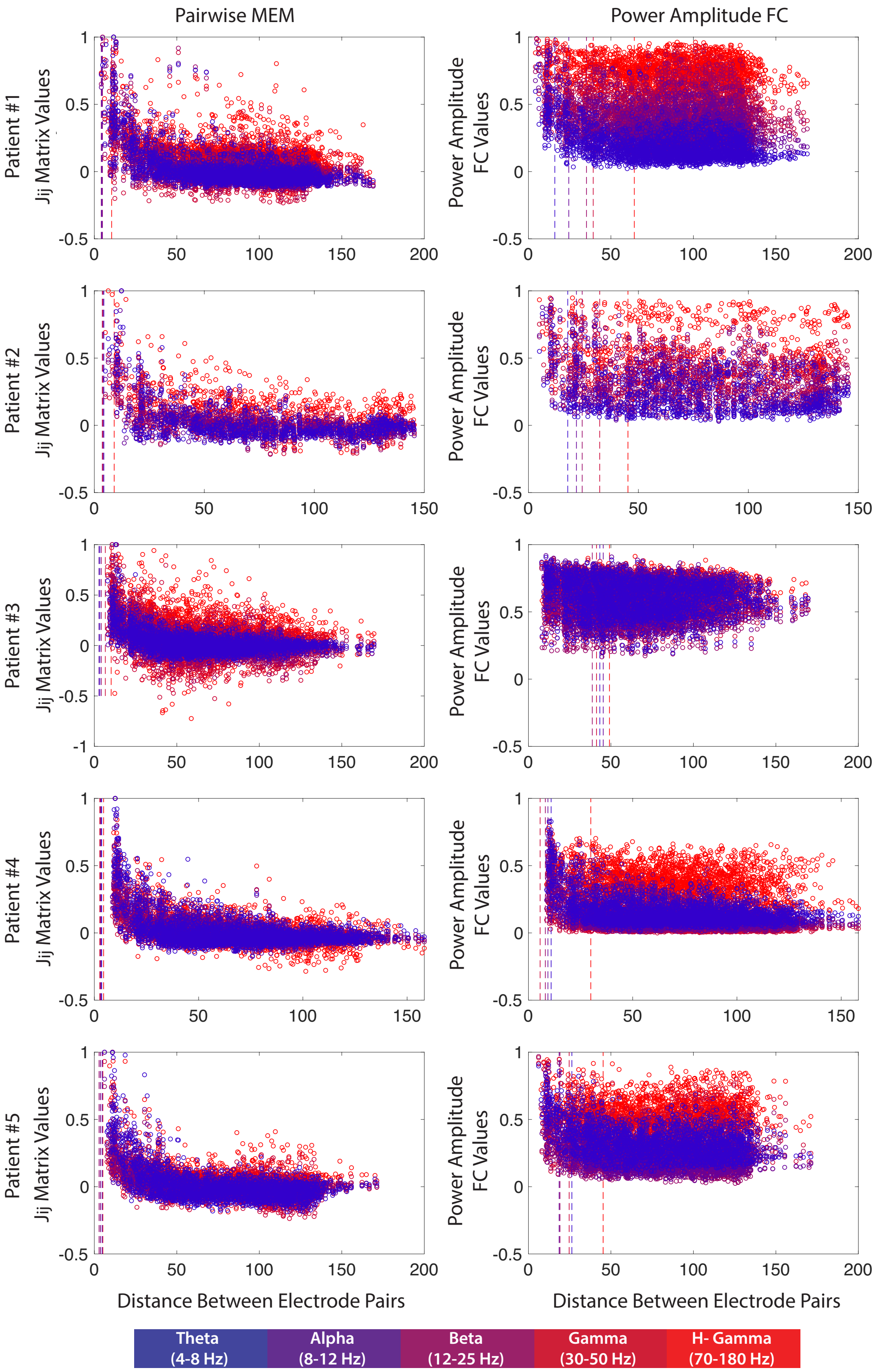


**Figure 2. Dependences of maximum entropy interactions and functional connectivity on inter-electrode distance.** (Left) We plot the normalized maximum entropy interactions $J_{ij}$ against the Euclidian distance between the corresponding electrodes $i$ and $j$. (Right) Similarly, we plot the normalized pairwise correlations in power amplitude activity (i.e., the functional connectivity) against the corresponding inter-electrode distances. Data are color-coded based on the frequency band. The dashed vertical lines indicate the average $J_{ij}$ and power FC values for each frequency band, weighted by the Euclidean distances between electrodes. Considering the distance-weighted average FC, we find that the distributions of FC become more skewed toward nearby electrodes as the frequency increases ($t$–test, $p < 0.05$ FDR). By contrast, the distributions of weighted average $J_{ij}$ are very similar and all are skewed towards nearby electrods. Although these weighted averages are very similar across frequency bands, we find that the distance-weighted normalized maximum entropy interactions $J_{ij}$ in the high $\gamma$ band are significantly higher across all patients than the interactions in other frequency bands ($t$–test, $p < 0.05$).

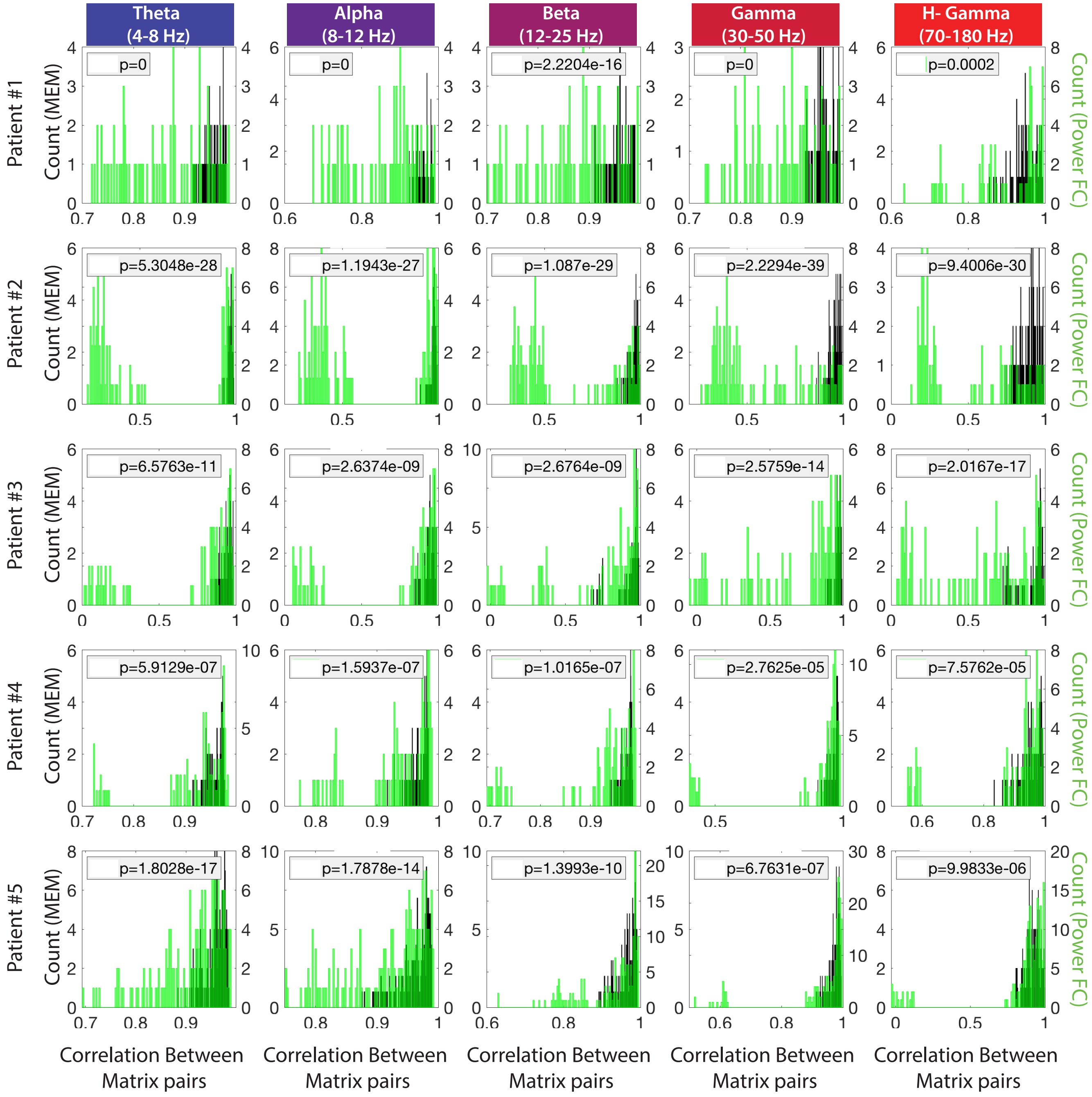


**Figure 3. Stationarity of maximum entropy interactions over time.** We calculate the correlations between the maximum entropy interactions $J_{ij}$ across all pairs of one-hour recordings (black). We compare these temporal correlations in $J_{ij}$ with the analogous correlations in the functional connectivity matrices (green). Each row represents results for one patient and columns are ordered based on frequency bands. We find that the pairwise maximum entropy interactions are significantly more consistent over time than the corresponding functional connectivities ($t$–test, $p < 0.05$).

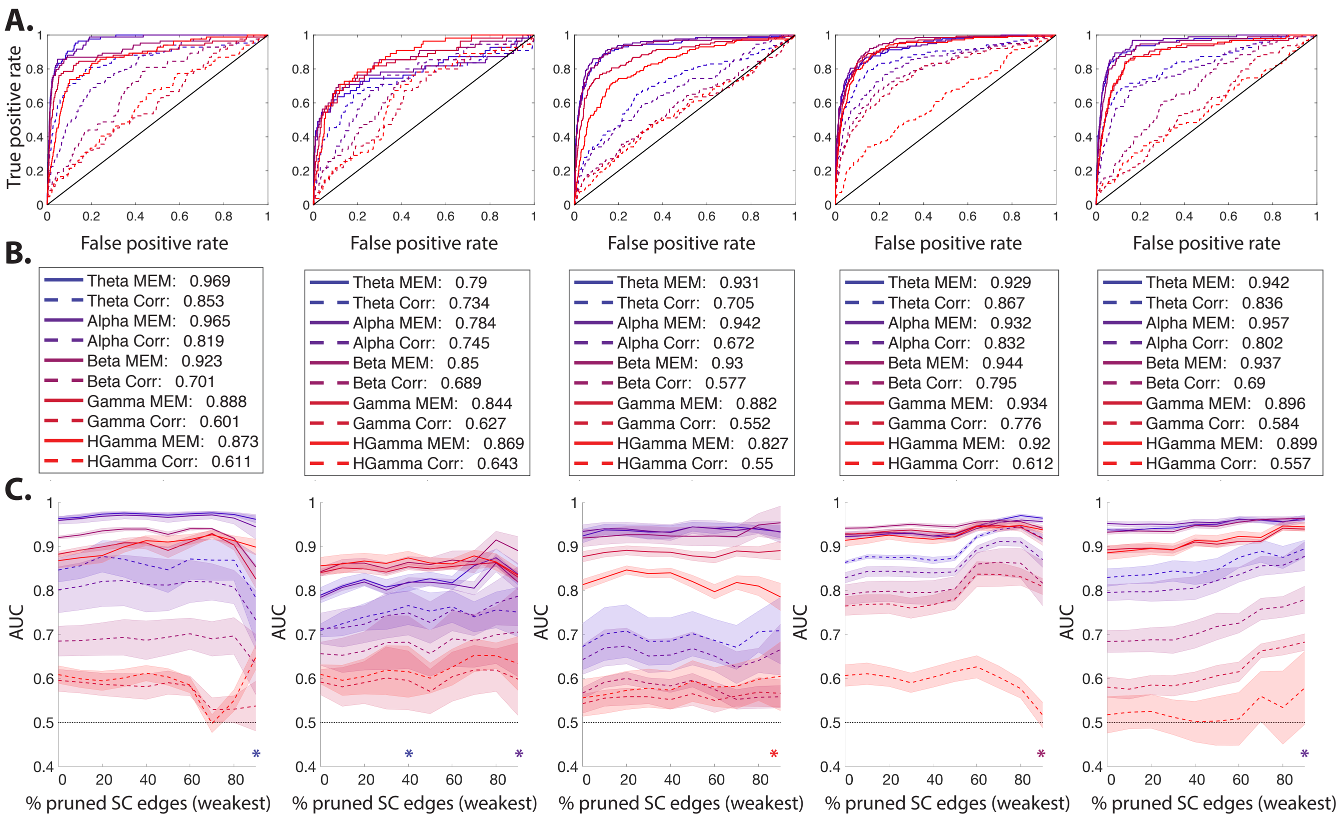


**Figure 4. Detection of structural connectivity from maximum entropy interactions and conventional functional connectivity.** ***(A)*** ROC curves for the identification of anatomically connected and unconnected ROI pairs (i.e., binarized SC) based on the average (over one-hour segments) maximum entropy interactions $J_{ij}$ (solid lines) and average time series functional connectivity (dashed lines) for all 5 patients (columns). Lines are color-coded based on frequency bands. ***(B)*** The AUC values for the curves in panel ***A*** ***(C)*** Comparison of the average AUC values (over one-hour segments) calculated using maximum entropy interactions (solid lines) and time series functional connectivity (dashed lines). The x-axis represents the percentage of the pruned weakest (i.e., low streamline count) SC edges. Similar to the results in Figure 3, the average AUC calculated from $J_{ij}$ interaction matrices are significantly (t–test, $p < 0.05$ FDR) higher than that of FC matrices across all frequency bands and across all pruning thresholds, with only a few exceptions (marked by '*').

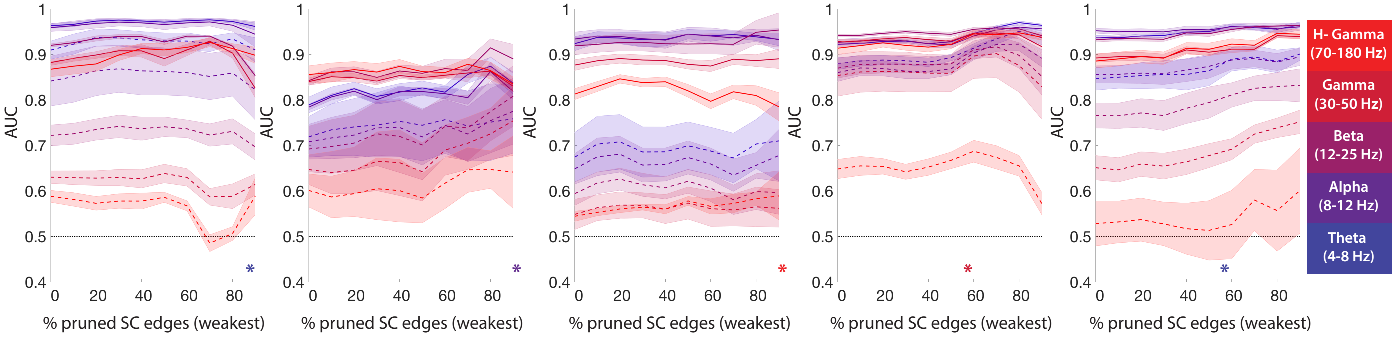


**Figure 5. Detection of structural connectivity from maximum entropy interactions and power amplitude functional connectivity.** We perform ROC analysis to quantify the abilities of maximum entropy interactions and functional connectivity to detect structural connections measured with DTI. The solid lines represent the mean AUC values (across all one-hour segments) calculated using the maximum entropy interactions for different SC pruning thresholds. Similarly, the dashed lines represent mean AUC values calculated using functional connectivity. Shaded regions represent standard deviations over the different one-hour segments. Each x-axis represents the percentage of the pruned weakest (i.e., low streamline count) SC edges, and the plots are color-coded based on frequency band. The average AUC values calculated from maximum entropy interactions are significantly higher than the corresponding values for power functional connectivity across all frequency bands and across all pruning thresholds (*t*–test, $p < 0.05$ FDR). These results hold for all patients with only a few exceptions (marked by '*').

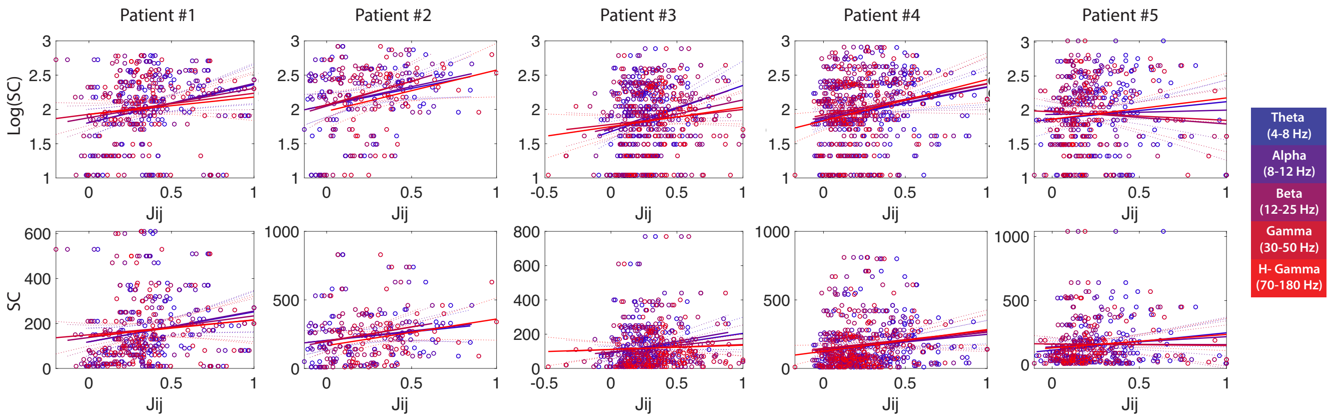


**Figure 6. Relationship between maximum entropy interactions and structural connectivity.** We plot the the normalized $J_{ij}$ interactions between anatomically connected brain regions against the weighted structural connectivities between the same regions. Structural connectivity weights represent log normalized fiber counts (top) and the raw fiber counts (bottom). Data are color-coded based on frequency band. Solid color-coded lines represent linear fits to the scatter plots for each frequency band and dashed lines indicate confidence intervals for the linear fits. Note that, except for patient #5 in the $\beta$ and $\gamma$ frequency bands, the structural connectivity strengths are positively correlated with the maximum entropy interactions $J_{ij}$.

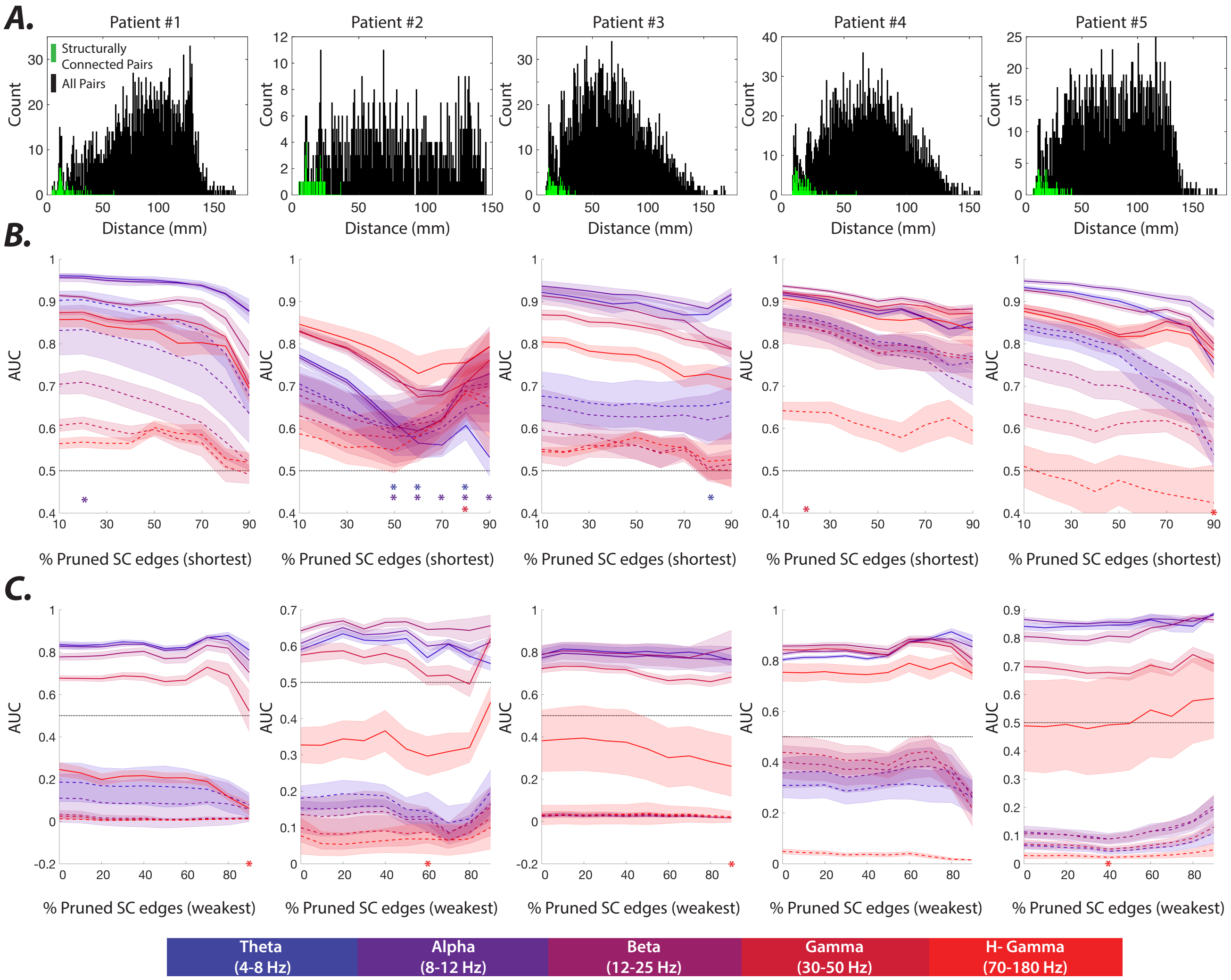


**Figure 7. The effects of inter-electrode distance on the detection of SC.** ***(A)*** Histograms of the distances between all electrode pairs (black) and those pairs that are structurally connected (green) for all 5 patients. Note that almost all structurally connected electrodes are proximal in our datasets. ***(B)*** We perform ROC analysis to test the abilities of the pairwise maximum entropy interactions (solid lines) and power amplitude functional connectivities (dashed lines) to detect the shortest structural connections. We plot the average AUC values versus the percentage of the shortest structural edges removed. Shaded regions represent the standard deviations over all one-hour segments of data. We find that the average AUC values calculated using $J_{ij}$ interactions are significantly ($t$–test, p $<$ 0.05 FDR) higher than those calculated using power amplitude FC across most frequency bands and most thresholds. There are only a few exceptions, the most pronounced of which occurred at lower frequencies in patient #2 (marked by '*'). ***(C)*** We now perform ROC analysis to detect structural connections using the maximum entropy interactions (solid lines) and power amplitude functional connectivities (dashed lines), but with the inter-electrode distances regressed out. Specifically, we regress out the proximity (i.e., $1/$distance) for each pair of electrodes in the $J_{ij}$ interaction matrices as well as the power amplitude FC matrices. We plot the averages (lines) and standard deviations (shaded regions) of the AUC values over all one-hour segments against the percentage of the pruned weakest (i.e., low streamline count) structural connections. Interestingly, even after accounting for inter-electrode distance, the maximum entropy interactions $J_{ij}$ identify structural connections across all frequency bands more reliably than the power amplitude functional connectivities ($t$–test, p $<$ 0.05 FDR); exceptions are marked by '*'. However, with the exception of patient #4, the $J_{ij}$ interactions for the high $\gamma$ band lose their ability to uncover the structural connectivity following the proximity regression, and we observe similar results for the power amplitude functional connectivities across all frequency bands. These results indicate that inter-electrode distance accounts for most of the observed correlation patterns in the high $\gamma$ band.

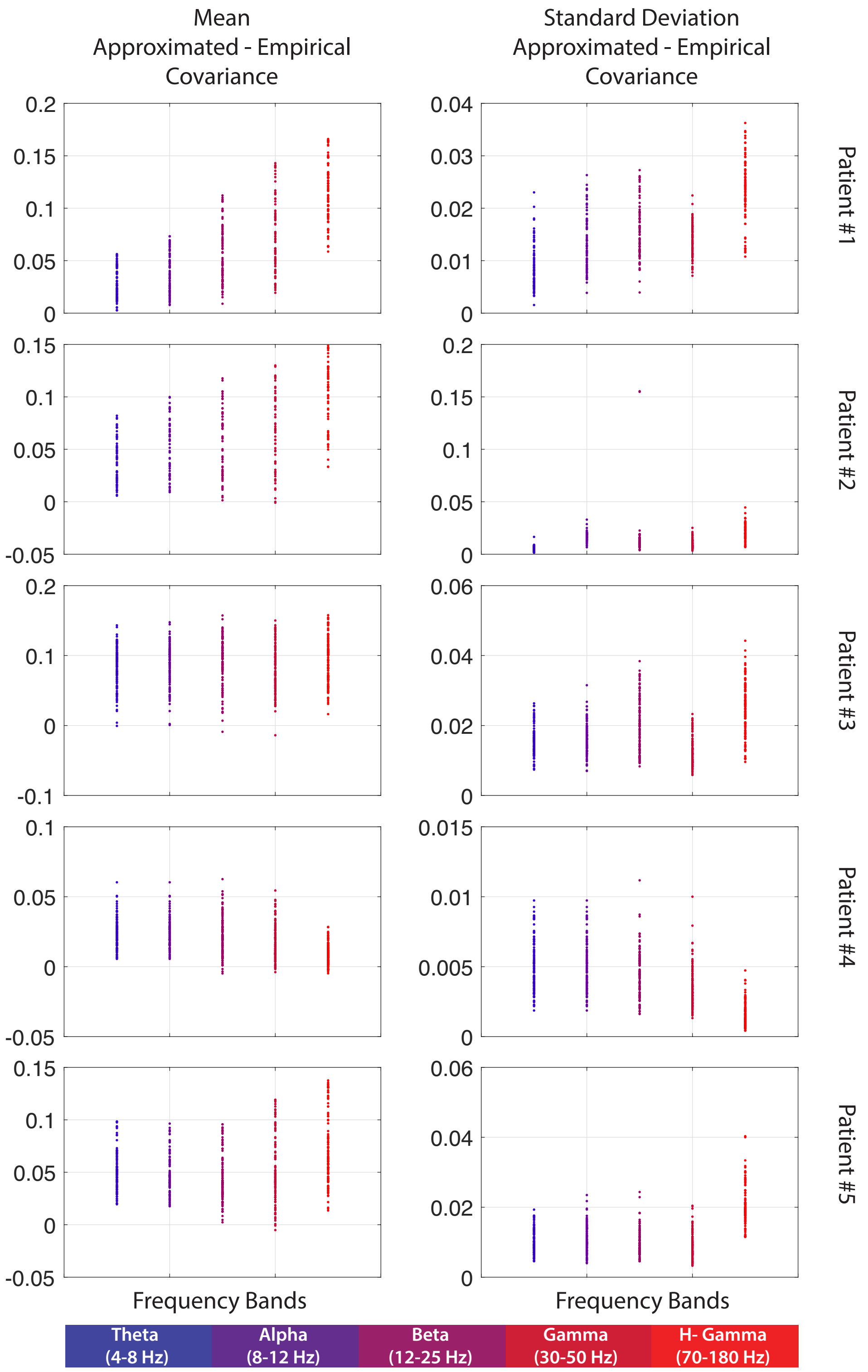


**Figure 8. Approximation errors for diagonal elements of the covariance matrices.** (Left) Average approximation errors for the diagonal elements of the covariance matrices across all patients and frequency bands. Each dot reflects the mean error of the approximated variance of a single electrode (i.e., a single diagonal element of the covariance matrix). Note that with the exception of a few instances across all electrodes, frequencies, and patients, the variance of the electrodes is on average significantly overestimated (paired $t$–test, $p < 0.001$ Bonferroni correction). (Right) Standard deviations of the approximation errors for diagonal entries of the covariance matrices. Throughout, dots are color-coded based on their frequency bands and each row displays the results for a given patient.

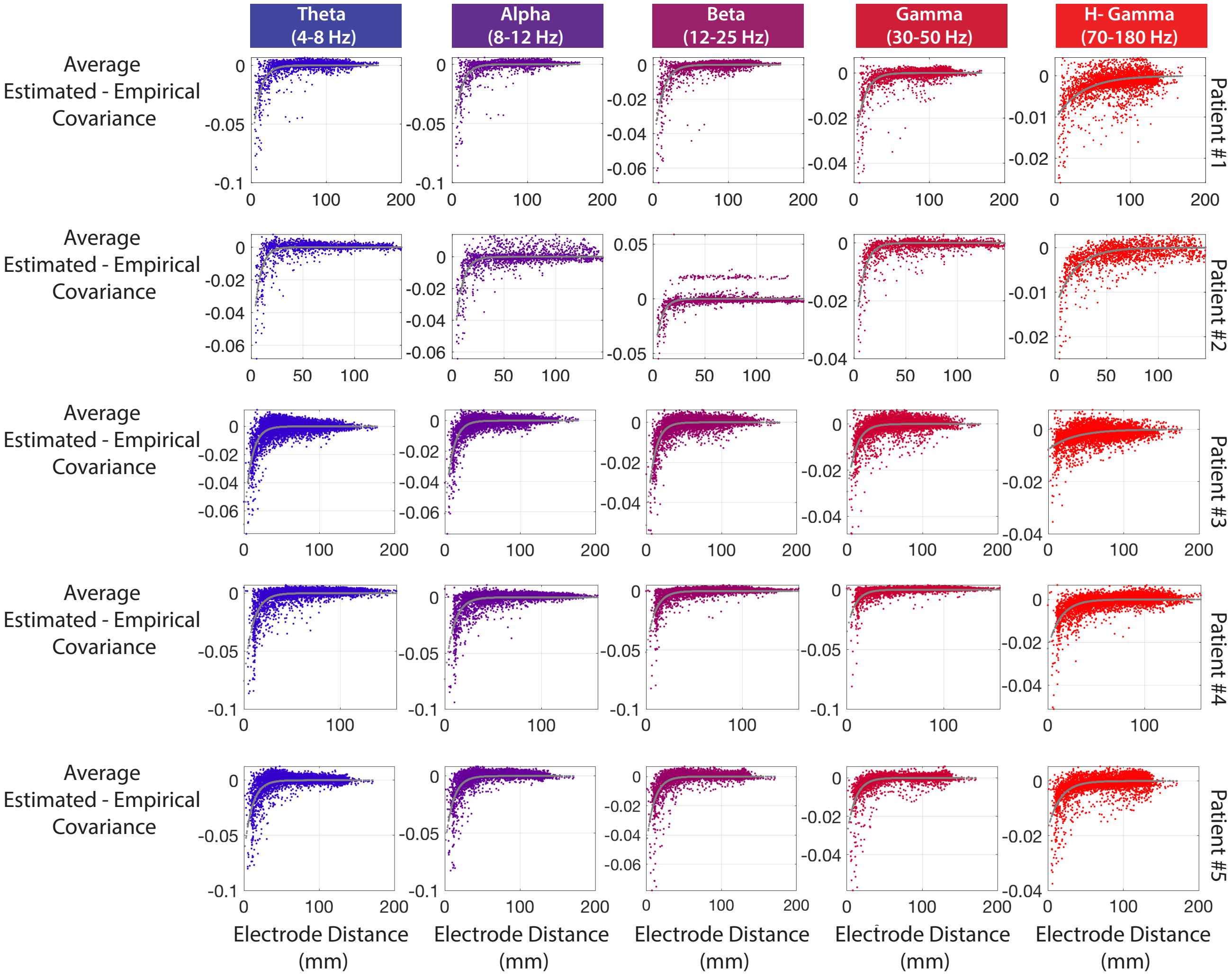


**Figure 9. Relationship between covariance approximation errors and the physical distances between electrodes.** Each row displays the relationship between average approximation error and inter-electrode distance for a given patient. Each column represents a specified frequency band. Gray lines indicate exponentials fit to the different scatter plots.

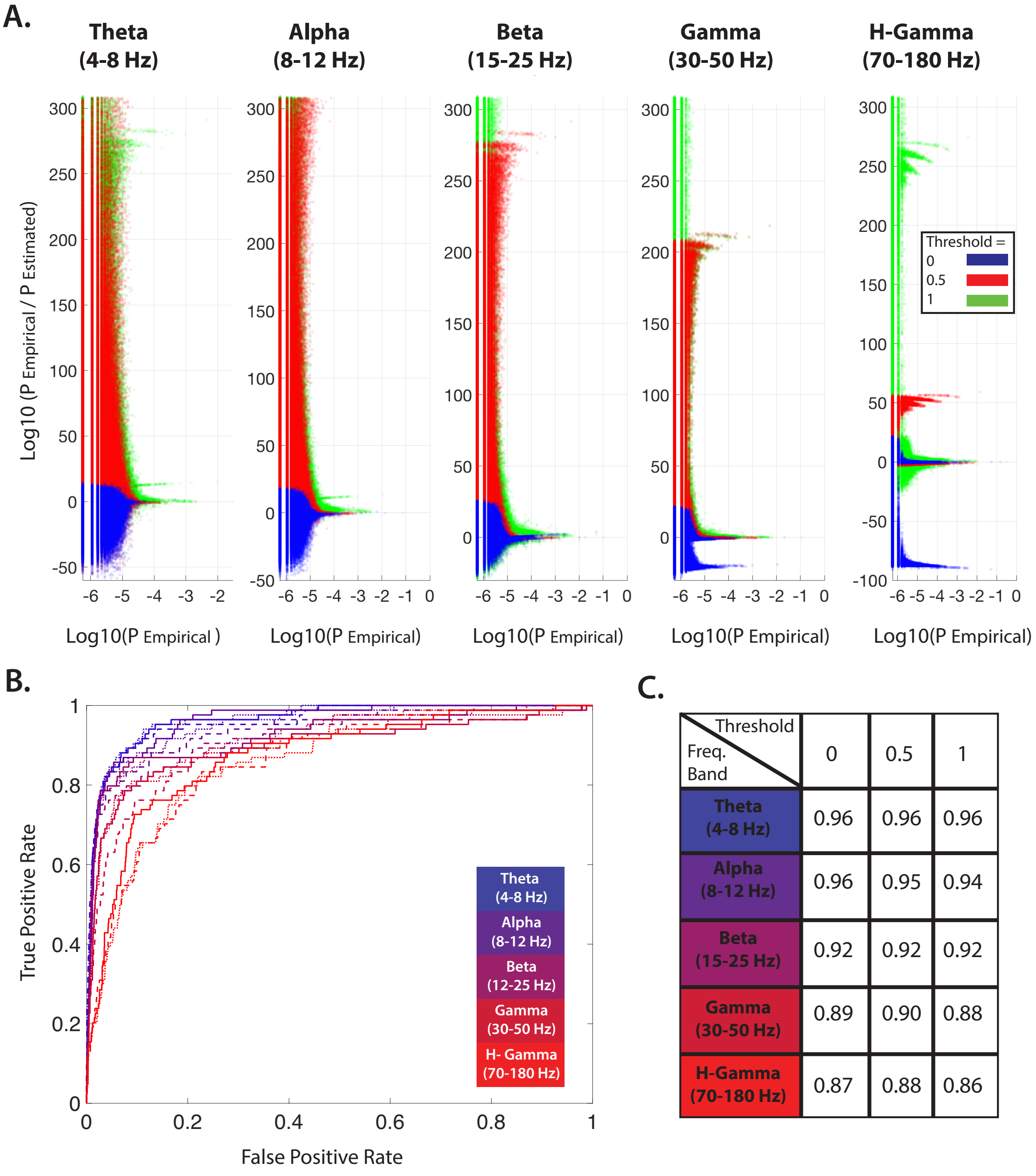


| Threshold / Freq. Band | 0 | 0.5 | 1 |
| --- | --- | --- | --- |
| Theta (4-8 Hz) | 0.96 | 0.96 | 0.96 |
| Alpha (8-12 Hz) | 0.96 | 0.95 | 0.94 |
| Beta (15-25 Hz) | 0.92 | 0.92 | 0.92 |
| Gamma (30-50 Hz) | 0.89 | 0.90 | 0.88 |
| H-Gamma (70-180 Hz) | 0.87 | 0.88 | 0.86 |

**Figure 10. Accuracy of the maximum entropy model in predicting activity patterns.** ***(A)*** Comparison of empirical and estimated (i.e., under the maximum entropy model) probabilities of observed activity patterns, where we show the data for patient #1 as an example. We plot the difference in log-likelihoods between the empirical and estimated probabilities versus the empirical log likelihood for each observed activity state, calculated for three different binarization thresholds: '0' (blue), '0.5' (red), and '1' (green). States with perfect predictions cluster along the line $x = 0$, while divergence from zero indicates estimation errors. The *partition function* (i.e., the normalization constant in the Boltzmann distribution) was approximated from the empirical probability of the *silent* state (see Ganmor et al. (2011); Tkačik et al. (2014) for more details). Note that the estimation error grows for states with lower empirical probabilities for all thresholds. Clusters of empirically observed states with notably higher predicted probabilities in $\gamma$ and high $\gamma$ bands (thresholded at '0') and lower predicted probabilities in the $\beta$ band (thresholded at '0.5' and '1') highlight the states where the model produces the largest errors. These results suggest that, although binarization thresholds affect the performance of the MEM and change the distribution of state estimation errors, the first and second-order interactions alone do not fully explain the probabilities of all observed states. ***(B)*** ROC curves for the identification of anatomically connected regions based on maximum entropy interactions calculated for different binarization thresholds: '0' (solid line), '0.5' (densely dashed line), and '1' (dashed line) for patient #1. Lines are color-coded based on frequency bands. Despite differences in state probability estimation errors of the pairwise MEM at different binarization thresholds, the ability of the estimated interaction matrices to predict the underlying SC remains virtually unchanged. The similar ROC curves and AUC values (presented in panel ***C***) highlight the robustness of SC detection to binarization threshold and model fit.

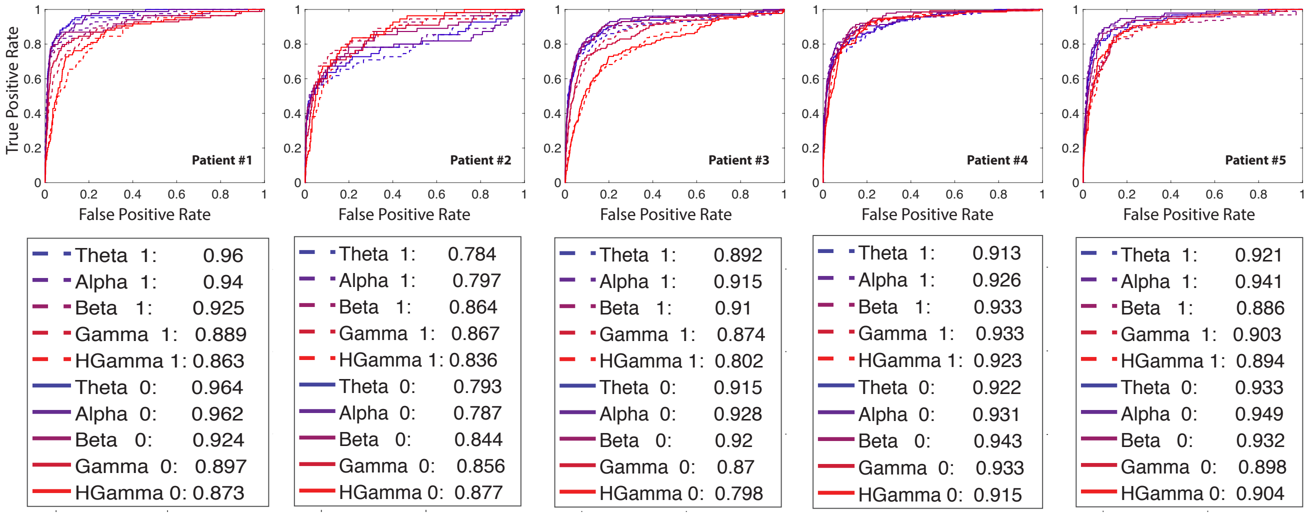


**Figure 11. Detecting anatomically connected regions with different binarization thresholds.** ROC curves for the identification of anatomically connected regions using the maximum entropy interactions for different binarization thresholds – ‘0’ (solid line) and ‘1’ (dashed line) – in a sample one-hour segment for all five patients. Lines are color-coded based on frequency bands. The AUC values are provided in the legend under each plot. The ability of the estimated interaction matrices to predict the underlying SC is robust to reasonable variation in the binarization threshold.

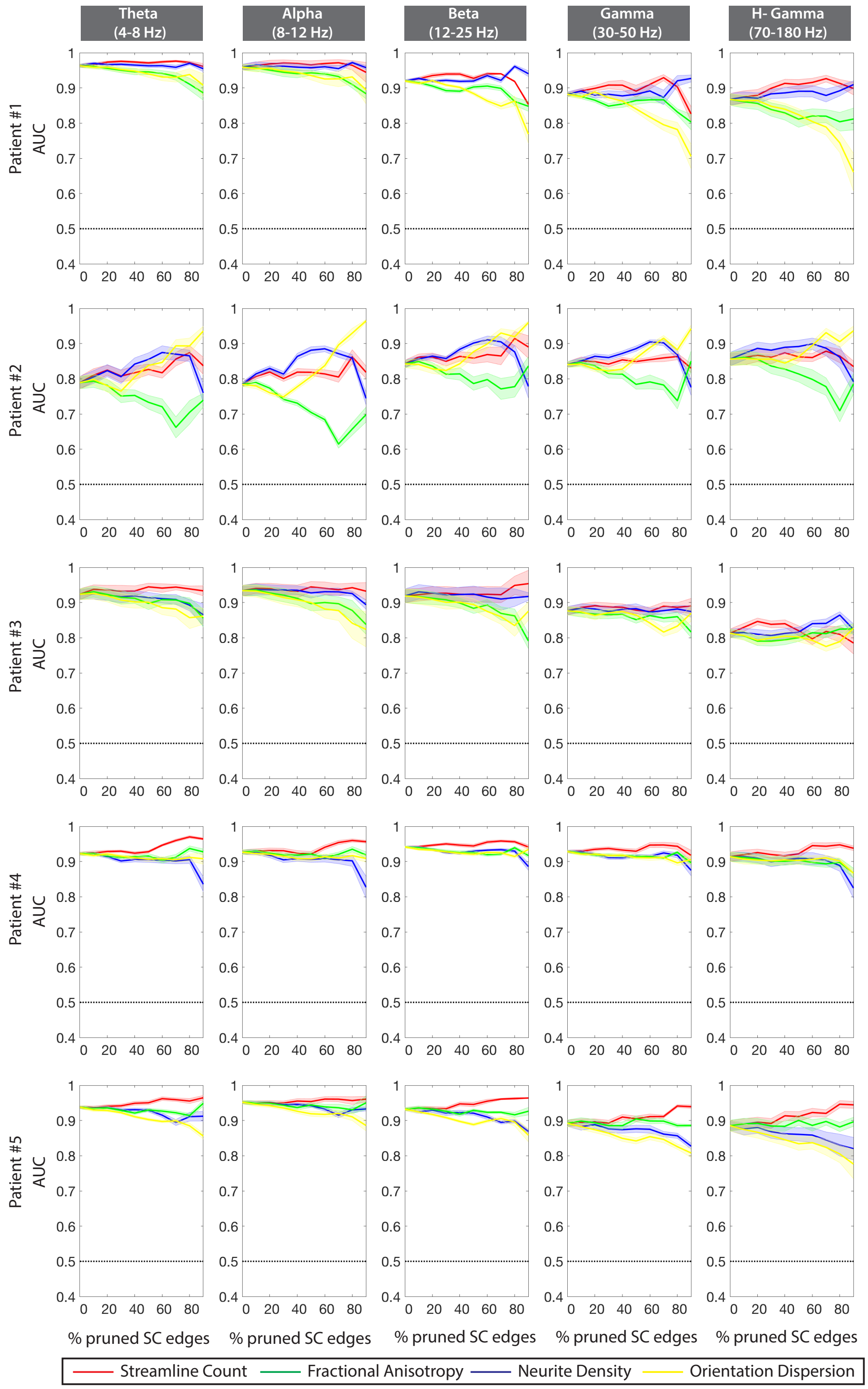


**Figure 12. Detecting different measures of structural connectivity.** The solid lines and shaded regions represent the mean and standard deviation (across all one-hour segments), respectively, of the AUC values calculated for different SC pruning thresholds using the maximum entropy interactions $J_{ij}$. The results are color-coded to reflect different measurements of anatomical connectivity between brain regions derived from DTI scans. The results for streamline count (SC), fractional anisotropy (FA) weighted, neurite density (ND) weighted, and orientation dispersion (OD) weighted adjacency matrices are represented by red, green, blue, and yellow lines, respectively. Each row displays the results for a given patient and each column displays the results for a given frequency band.
